# “Quantitative analysis reveals the basic behavioural repertoire of the urochordate *Ciona intestinalis”*

**DOI:** 10.1101/382465

**Authors:** Jerneja Rudolf, Daniel Dondorp, Louise Canon, Sonia Tieo, Marios Chatzigeorgiou

## Abstract

Quantitative analysis of animal behaviour in model organisms is becoming an increasingly essential approach for tackling the great challenge of understanding how activity in the brain gives rise to behaviour. In addition, behavioural analysis can provide insight on the molecular basis of nervous system development and function as demonstrated by genetic screens focused on behavioural phenotyping in some genetically tractable model organisms. The progress in building low-cost automated tracking setups, together with advances in computer vision machine learning have expanded the repertoire of organisms which are amenable to quantitative behavioural analysis. Here we used automated image-based tracking to extract behavioural features from an organism of great importance in understanding the evolution of chordates, the free swimming larval form of the tunicate *Ciona intestinalis* which has a compact and fully mapped nervous system composed of only 231 neurons. We analysed hundreds of videos of larvae and we extracted basic geometric and physical descriptors of larval behaviour. Most importantly, we used machine learning methods to create an objective ontology of behaviours for *C. intestinalis* larvae. We identified eleven behavioural modes using agglomerative clustering. This approach enabled us to produce a quantitative description of the basic larval behavioural repertoire. Furthermore, we tested the robustness of this repertoire by comparing different rearing conditions and ages. Using our pipeline for quantitative behavioural analysis, we successfully reproduced the known photoresponsive behaviour and the first demonstration to our knowledge that *C. intestinalis* larvae exhibit sensory arousal and thigmotaxis, both of which can be modulated by the anxiotropic drug modafinil. Remarkably, by comparing the behaviour between animals assayed individually or in small groups, we found that crowd size influences larval behaviour. This study shows that *C. intestinalis* larval behaviour can be broken down to a set of stereotyped behaviours that are used to different extents in a context-dependent manner. Furthermore, it raises exciting possibilities such as mapping behaviour to specific neurons of this compact chordate nervous system and it paves the way for comparative quantitative behavioural studies as a means to reconstruct the evolution of behaviour, especially in the chordate lineage.

## Introduction

Close observation of living animals can reveal the large repertoire of behaviours they use to interact with the world. Animals can crawl, swim, run and fly to move from one place to another. Many animals perform extremely complex behaviours to attract mates, exhibit parental care and establish their position in social hierarchy. Numerous species are able to build elaborate structures, ranging from spider webs for catching preys to bird nests for shelter and raising of offspring. Some can even make and operate tools. These observations have led to two important challenges for the scientific community to pursue.

The first is to obtain a detailed understanding of how nervous systems generate behaviour. Modern approaches to tackle the first challenge include techniques for recording and targeted manipulation of neuronal activity using a wealth of molecular and cell type information (1). However, to fully understand the function of neural circuits, we need to obtain an equally precise and detailed understanding of behaviour (2, 3).

Behaviour is a process that is characterised by dynamic changes, and complex sequences of events that are often convoluted with noise. Therefore, measuring animal behaviour using manual approaches can be time consuming, and prone to errors, the latter especially in cases where a behavioural event is taking place over a very short or very long time scale, making it difficult to be detected by the experimenter. Modern computational analysis methods and accessible hardware for recording videos with high temporal resolution make it possible to observe and quantify behaviour in a more comprehensive, accurate and automated approach (4-7).

Automated behavioural analysis has been used to divide and classify behaviour into distinct modules, and has been extensively demonstrated in several organisms, including worms (8, 9), flies(10), zebrafish (11, 12) and mice (13). Despite the morphological and locomotor differences between these organisms, automated tracking systems coupled to machine learning can transform what appears as complex behaviours into a sequence of more basic motor patterns that are executed in a particular frequency and order. Overall, this approach can result in a simplified description of how behaviour is organised and carried out.

The second challenge is to understand how behaviours and nervous systems co-evolved through time. Most efforts to achieve this goal have focused on the dissection of the developmental and genetic mechanisms driving the evolution of neuronal circuits linked to species-specific behaviours (14, 15). The advent of automated tracking and analysis methods that allow for the recognition and segmentation of morphologically diverse organisms, offer the opportunity to perform quantitative behavioural analysis of key organisms across different taxa (16-18). This approach could provide important insight into conserved, as well as novel behavioural programs and how they are linked to major evolutionary transitions in nervous system structure and cell type composition.

In this study, we used the tunicate *Ciona intestinalis* to tackle these two challenges. Tunicates are the closest relatives of vertebrates and have been successfully opted as models to study the evolution of chordates(19). They have larvae with a chordate body plan and development(20). The two most intensively studied tunicate species are *Ciona intestinalis* and *Ciona robusta. Ciona intestinalis* features a defined cell lineage, an extensive genetic toolkit and a sequenced genome that shares a high number of homologous genes to its vertebrate counterparts. It has been extensively used in studying the chordate origins of many biological processes and has been particularly successful as a model for studying the development and evolution of the chordate nervous system (21). It possesses a dorsal central nervous system for which a documented synaptic connectome of the 177 CNS neurons has recently been published. Together with a publication of the peripheral nervous system connectome demonstrating the presence of an additional 54 neurons, these studies have laid out the foundations for future functional studies (22-24). This makes *C. intestinalis* the second organism with a complete connectome available after *Caenorhabditis elegans(25).*

An adult *C. intestinalis* animal can release hundreds of eggs together with sperm. Gametes undergo fertilization and through a series of stereotyped developmental steps, with the embryos taking the form of hatching lecithotrophic larvae. When the larvae hatch from the chorion, they find themselves in the water column. Following the hatching event larvae swim upwards towards the water surface by negative gravitotaxis using the otolith cell. Ablation experiments have shown that animals lacking the ocellus are also capable of this behaviour, indicating that the ocellus is not involved in gravitotaxis.(26, 27). Later on, larvae exhibit negative phototaxis, swimming away from the bright surface to deeper waters in a behaviour that possibly aims to identify suitable substrates for settlement (28, 29). The swimming larvae are thought to display three types of swimming activity: tail flicks, spontaneous swimming and shadow response(30). Larvae under constant illumination swim more frequently and for more extensive periods of time earlier in life, up to 2 hours post hatching. A behaviour that develops later in development is the shadow response, where dimming of light results in symmetrical swimming. After two hours post hatching the tail beating frequency increases(30). *C. intestinalis* larvae have also been shown to exhibit both sensitization and habituation to light (31, 32). A recent study reported that distinct groups of photoreceptors mediate negative phototaxis and dimming behaviour in *C. intestinalis* larvae(33). This functional insight was achieved through behavioural analysis on mutant lines, demonstrating the power of genetics in combination with behavioural analysis and knowledge of the larval connectome. However, this is not the first attempt to link behaviour to its genetic underpinnings in *C. intestinalis* as it has been shown that targeted knockdown of opsin1 results in a loss of the light OFF response in larvae(34). Beyond phototactic and gravitotactic behaviours there is evidence that hints to the possibility that the larvae can exhibit chemotactic(35) and mechanosensory behaviours(36-40). However, these have remained largely unexplored.

Over the past decades, the biological research community has used *C. intestinalis* as a model for uncovering the basic principles of chordate embryonic and post-embryonic nervous system development. We believe that by building on this work (e.g. taking advantage of the numerous characterised promoters, known cell lineage etc), we can use the compact nervous system of the larva in order to uncover the basic molecular and cellular mechanisms underlying chordate nervous system function. We have the tools required to deconstruct the molecular and cellular components of the larval nervous system(41) and the opportunity to adopt existing toolkits to probe its functions(42). Here, we attempt to match the growing insights into the structure and function of the *C. intestinalis* nervous system with a detailed characterization of its behavioural output. To achieve this, we collected a large dataset of recordings of free-swimming *C. intestinalis* larvae on a custom IR-illuminated and temperature-controlled set-up. We analysed the recordings using ToxTrac(43) and custom written Python scripts to obtain positional data and descriptors like speed and its variability, turning angles, and path complexity. To describe behaviour both accurately and objectively, we used unsupervised clustering methods to identify distinct behavioural clusters in our dataset, ultimately arriving at a quantifiable distribution of detectable behavioural components. This approach enabled us to study how this distribution of behavioural components changed under the influence of different rearing conditions, at different ages and in response to different sensory cues. Strikingly, we show that *C. intestinalis* larvae exhibited a state of sensory arousal when first transferred into the behavioural arena. This sensory arousal state and thigmotaxis behaviour could be modulated by the anxiotropic drug modafinil.

## Results

### Defining behavioural parameters and modes

In order to probe the behavioural dynamics of *C. intestinalis* larvae, we built an inexpensive tracker, with temperature control and the ability to deliver light stimuli with accurate spatiotemporal control. To maximize the open-source accessibility and modularity of the trackers we used Arduino based circuits for controlling the various tracker components and 3D printed housing for the agarose plated behavioural arenas. The agarose arena was housed inside a 3D printed ring fitted with infrared (IR) LEDs (Fig 1A).IR LEDs were used in order to visualise the animals in a dark-field configuration, while at the same time avoiding stimulating their photoreceptors with visible light, which they can detect and modify their behaviour (31-33). We imaged approximately 550 *C. intestinalis* animals behaving freely at two different frame rates: During the first 15 minutes after transferring animals to the setup we imaged at 10 frames/s and subsequently we imaged the same animals for 5 minutes at 30 frames/s. Following acquisition and processing, the videos were analysed on the tracking program ToxTrac(43) (Fig 1C, D). Due to the lack of previous attempts to track *C. intestinalis* larvae with automated image tracking software, we tried a series of available programs. We found that ToxTrac worked best in our hands especially because it did not require specific knowledge of the animal’s body geometry and because it was particularly robust against false positives and identity switches. The position of the animal was tracked by the centre of its detected shape, as identified by ToxTrac. From all the videos tracked we obtained around 850 traces for further analysis and quantification.

Below we introduce some of the descriptive parameters used and the reasoning behind their definition.

*C. intestinalis* larvae have been reported to modulate their locomotor activity levels through “bursts” of spontaneous activity (26, 30, 44). We decided to introduce a parameter that would provide a quantitative descriptor of locomotory activity, termed Activity coefficient (AC) and defined as the fraction of time an animal spent locomoting. Filtered speed values of 200 μm/s and above were considered as active, which in practical terms includes all actively swimming animals as well as movement of the animals’ centre-point due to tail flicks and twitching.

Planktonic larvae from different species exhibit a wide range of paths during locomotion(45), often dictated by their biomechanics (44, 46-48). Notably, reduced food availability can result in nonlinear paths, while directionality in trajectories can often be indicative of behavioural responses to environmental stimuli (49, 50). While recording animals in the behavioural arenas, we observed that they exhibited a large repertoire of trajectories while swimming in the arena. Some of the animals were swimming in straight lines with relatively few turns and changes in speed, representing what an observer would describe qualitatively as low complexity trajectories. In contrast, other animals moved in a more “complex” fashion, exhibiting sudden changes in direction and moving in spiral or circular paths (see Fig 1E for example trajectories). We decided to quantify the local path complexity using a method presented by Roberts et al. (51). In summary, this method uses embedding matrixes for positions in a specific time window, over which the local path complexity is calculated in bits of entropy. Minimal complexity values calculated by this method correspond to the most predictable trajectory or, in other words, the most invariable movement in terms of represented speeds and directions in the time window (see Fig 1F for example traces with decreasing local complexity).

While observing the animal trajectories, we noticed that a significant fraction of animals swam near the edges of the arena. This behaviour has been previously identified in other organisms (13, 52-54) including humans(55-57) and has been termed thigmotaxis. We decided to quantify the thigmotactic behaviour of the larvae. To define thigmotaxis, we divided the arena into two concentric zones of equal surface area. Animals in the outer zone were considered thigmotactic (Fig 1B). The amount of thigmotaxis is quantified in two measures: “Total Time spent in Outer zone” (TTO) and “Total Distance travelled in Outer zone” (TDO), as utilised for zebrafish previously (58).

A long-term goal of quantitative animal behaviour analysis is to identify the behavioural strategies that animals use to solve tasks and to dissect the underlying neuronal mechanisms. One approach to reach this goal is to generate a structured quantitative description of behaviour, by identifying stereotyped behavioural components. In order to identify these behavioural components in behaving *C. intestinalis* larvae, we used unsupervised clustering methods on a small feature-set (ρ-speed, Δρ-acceleration and Δϕ-turn angle values; see methods section for details), and we were able to identify and classify 11 distinct clusters in our data set. We assigned a relevant descriptor of the behavioural mode to each cluster according to its biological interpretation. This was done by inspecting the original videos and superimposing corresponding cluster data. When unable to resolve a biological difference between two clusters, we assigned them the same descriptive name. We list the behavioural modes in approximate order of speed of movement in Table1 below (see Fig 1G for some example traces of different modes and S1-S5 Video for example videos). This simple ontology of the objectively detectable behavioural modes in our data set served as an extra tool in assessing the effects of different stimuli and genetic or pharmacological perturbations on the behavioural repertoire of the larvae.

**Table 1:** Behavioural clusters with names.

| Cluster | Descriptive name |
| --- | --- |
| 01 | Inactive 1 |
| 02 | Inactive 2 |
| 03 | Small twitches |
| 04 | Large twitches |
| 05 | Collision or deceleration event |
| 06 | Mode change 1 |
| 07 | Mode change 2 |
| 08 | Slow active swimming |
| 09 | Medium active swimming |
| 10 | Fast swimming 1 |
| 11 | Fast swimming 2 |

**Fig 1.**
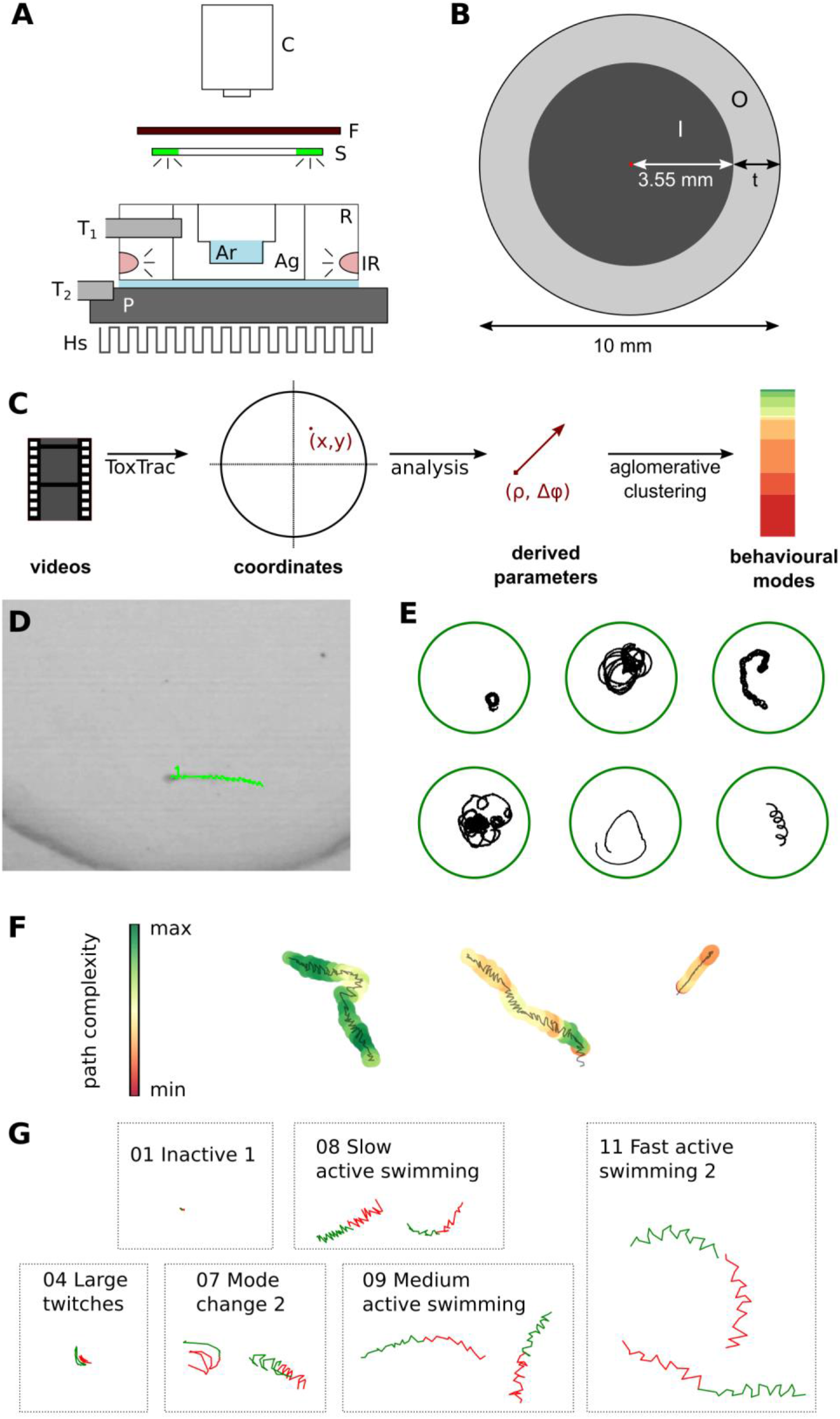
Setup and analysis methods. (A) Setup: C- camera, F- IR filter, S – ring with stimulation LED-s, T1 – Thermometer 1 measuring local temperature in the agarose, T2 – Thermometer 2 measuring cooling plate temperature, P – cooling plate, Hs – heat sink, R – PLA ring holding the arena, Ag– agarose, Ar– Arena with animals, IR – IR illumination. (B) Arena dimensions and areas used in thigmotaxis measures I-inner arena with radius 3.55 mm and O– outer thigmotaxis zone; with t= 1.45 mm the zones have equal surface area and the width of the thigmotaxis zone is above one animal body-length. (C) Workflow chart. (D) An example animal being tracked with the ToxTrac software (E) Some example trajectories of wild type animals swimming for 5 minutes (F) examples of different local complexity of a trace - each trajectory is coloured by local complexity which is calculated over a 3 s window and the total span of each trace is 6s (G) Examples of traces spanning 50 frames based on which the current behavioural mode was calculated. In green are the 25 frames before the current time-point and in red the 25 frames later.

### Adaptation to the arena

The introduction of an animal to a new environment, such as a tracking arena, is one of several potential triggers of generalised nervous system arousal (59-61). Arousal can also be observed in response to stimuli associated with harm, such as strong mechanical stimuli. In the case of the nematode *C. elegans,* transferring an animal from one plate to another using a metal pick can result in a temporally defined state of arousal demarked by higher motor activity that eventually returns to baseline roaming locomotor activity(62). A similar observation has been made with mice when placed in an open-field arena(63). This period is often termed the adaptation period. Given that arousal mechanisms are evolutionarily conserved (64), we asked whether *C. intestinalis* larvae were subject to generalised nervous system arousal as a result of the transferring process to the tracking arena and the exposure to a new environment.

In order to answer this question, we decided that before recording the videos used for analysing baseline behaviour, each animal would first be recorded for a 15 min period (Fig 2). In the first minutes of the animal being exposed to the new environment its speed was generally higher (see Fig 2A for example traces), which we quantified as the slope of linear regression over the average speed values of around 100 animals (Fig 2B, C). From these results, we inferred that the animals adapted to the arena within approximately 6 minutes. We compared some basic behavioural parameters of individual animals between the first, second and last third of the 15 min adaptation period (Fig 2 D-G). There are significant differences in both median speed (Fig 2D, median at 267 μm/s in the initial 5 min vs. 101 μm/s in the second and 113 μm/s in the last 5 minutes; *p* k 0.001 for both comparisons) and maximum speed of the animals (Fig 2E, median at 1310 μm/s in the initial 5 min vs. 1003 μm/s in the second and 795 μm/s in the last 5 minutes; *p(1 vs. 2)* = 0.0013, *p(1 vs. 3)* < 0.001, *p(2 vs. 3)* = 0.0 48). Interestingly, as a result of sensory arousal, *C. intestinalis* larvae exhibited higher minimum path complexity in the first 5 minutes after they are placed in the arena (Fig 2F, 4.349 bit in the initial 5 min vs 4.297 and 4.293; *p(1. vs. 2.)* = 0.027, *p(.1 vs. 3.)* = 0.0038). This observation suggested that the animals followed a more unpredictable, or chaotic trajectory during the original arousal period.

**Fig 2.**
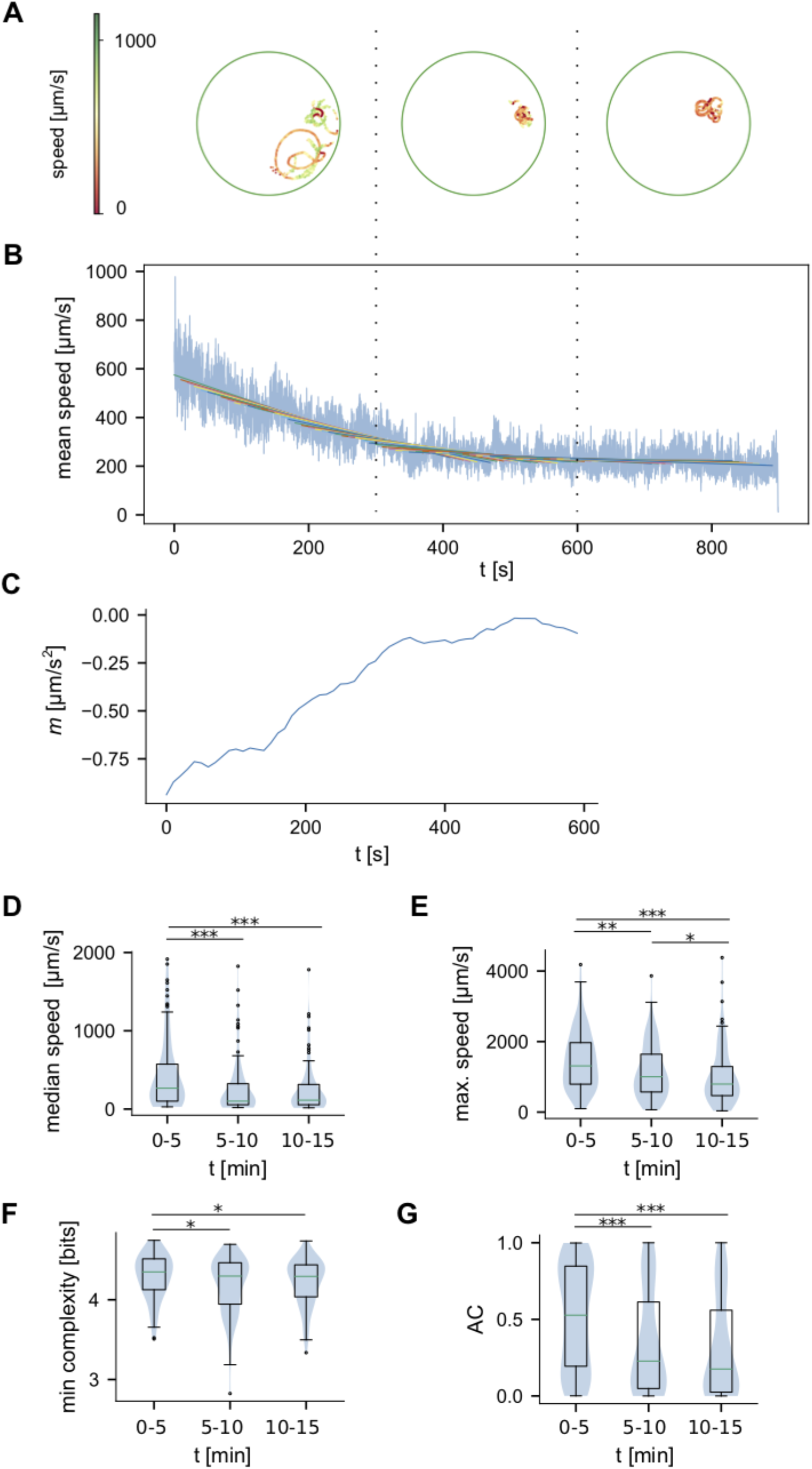
Adaptation of animals to the arena. Analysis of adaptation behaviour of larvae in the first 15 minutes after introduction to the arena. (A) An example trajectory of a path during the adaptation period. The path is plotted in thirds each corresponding to 5 minutes of the recording as illustrated by the dotted lines linking this and the next panel (B) Average speed of 105 animals during the 15 minutes after introduction to the arena shows higher speeds immediately after the animals were exposed to the new environment. (C) Slope of linear regression fitted to the average speed (window 5 min, step size 10 s) shows the stabilisation of speed after ca 6 min. We compared (D) median speed, (E) maximum speed, (F) minimal reached path complexity and (G) activity coefficient between the first, second and last third of the 15 min adaptation period (N_1_= 105, N_2_=102, N_3_=92).

As they adapted they followed trajectories exhibiting lower entropy and thus higher degree of predictability. In other organisms such as birds the trajectory component showing higher entropy is thought to be associated with navigational uncertainty(65). Notably, the activity coefficient (Fig 2G, median at 0.53 vs. 0.23 and 0.18; *p* < 0.001 for both comparisons) of the aroused animals is also significantly higher in the first 5 minutes, suggesting that bouts of inactivity are suppressed during this period. Finally, unlike mice in open-field tests, which show high thigmotaxis during the initial period(63), we found no significant changes in thigmotaxis in the case of aroused larvae vs. animals that have adapted (data not shown).

### Behavioural effects of larval crowd size

In a laboratory setting we often study behaviour using individual animals in isolation. However, in the wild, organisms are hardly ever acting in absolute isolation. Across the animal kingdom, locomotion and response to environmental stimuli are influenced by both direct and indirect interactions with conspecifics (66, 67). For example, interactions between individuals are important in mating(68), in the protection of colonies from pathogens(69), and in response to predators(70, 71). Previously it has been shown that mechanosensory interactions between desert locusts can lead to a dramatic phenotypic change where the animals switch from a cryptic solitary phase to a gregarious phase(72). More recently, in the genetically tractable organism *Drosophila melanogaster* it has been shown that mechanosensory interactions between adults drive collective response towards a sensory cue(73).

The behaviour of *C. intestinalis* larvae in large groups has been studied to some extent, e.g. the change in distribution because of water agitation(29) or stimulation with light (30). In our study, we focused on the possibility of behavioural effects of small group interactions. To achieve this, we compared animals that were alone in the arena with animals recorded in pairs or groups of three. We found small differences in the basic behavioural parameters (Fig 3). Crowd size did not affect median (Fig 3A) or maximum speeds of movement significantly, but had a weak effect on the path complexity and activity coefficient (Fig 3B, C). The average path complexity was lowest for animals in the crowd size 3 group (6.281 bits, compared to 6.290 bits for animals in crowd size 1 group, p=0.05) and so was the median AC value (0.63 compared to 0.82 and 0.75 for animals in crowd size 1 and 2 groups respectively, *p(1. vs. 3.)* =0.045). We also found slightly more animals with higher thigmotaxis values, with the difference only significant for TDO (Fig 3D) between crowd size 1 and 3 (median TDO is 0.17 for crowd size 1 vs. 0.47 for crowd size 3, p= 0.0227). The distribution of different behavioural modes shows higher representation of the less active modes corresponding to more time spent inactively in crowd size 3 animals (Fig 3E). In Fig 3F we present sample ethograms of 6 individual animals in different crowd size experiments. The smaller representation of higher speeds was also apparent when we plotted the distribution of speed values with the corresponding turn values on a polar scatterplot, presented for the different crowd sizes in Fig 3G.

**Fig 3.**
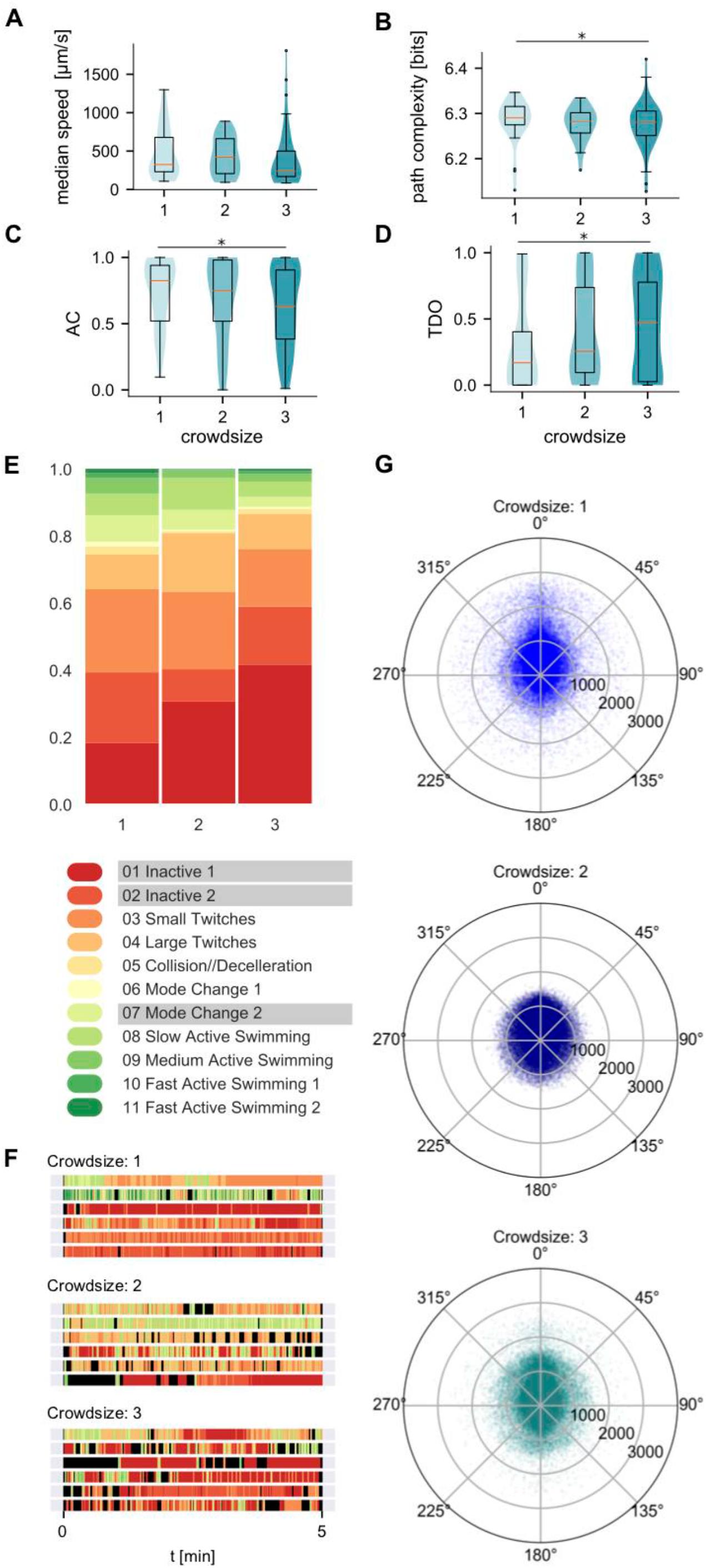
Crowdsize effects on behaviour. Crowdsize effects on (A) median speed, (B) path complexity, (C) activity coefficient and (D) TDO. (E) Distribution of behavioural modes in animals in experiments with different crowdsizes. The modes underlined in the legend are the biggest contributors to the Chi2 statistics (data in S1 Table) (F) Example ethograms of individual animals in crowdsize experiments. Each line represents a 5 min recording and is coloured based on the assigned behavioural mode, black colour represents missing frames where modes could not be assigned. (G) Polar scatterplots of filtered speed values vs turn values for different crowdsizes (N1=33, N2=22, N3=46; number of points per polar plot is 100000).

### Age related changes in behaviour

Behavioural changes that result from aging have long been the focus of various senescence studies, but there are similarly important age related changes underlying the normal development of the animals’ behaviouralrepertoire through time(74-79). For larval *C. intestinalis,* light and gravity dependent behaviours have already been shown to change with age (26, 31). Here we examined the development of baseline behaviour in the early hours post-hatching. In animals reared and recorded at 14 °C, we observed that in the first hour after hatching they were mostly inactive, followed by a period of time when the larvae twitched very actively and flicked their tails but would not cover a lot of distance swimming. In Fig 4 we show some of the basic behavioural descriptors compared between ages of WT animals reared at 14°C. Both maximum and median speeds were lowest for animals immediately after hatching (Fig 4A, B), accompanied by very low AC (Fig 4C). While animals 1h post fertilisation already achieved higher median speeds and generally had a very high AC, their movement was less directional as can be seen by the distribution of turn values versus speed values (Fig 4E) and the high representation of twitching modes in their behavioural repertoire (Fig 4D). To minimise any potential skewing of the data because of age-dependent changes, we used animals of 2–8 hours post hatching age for all later comparisons, unless specified otherwise.

**Fig 4.**
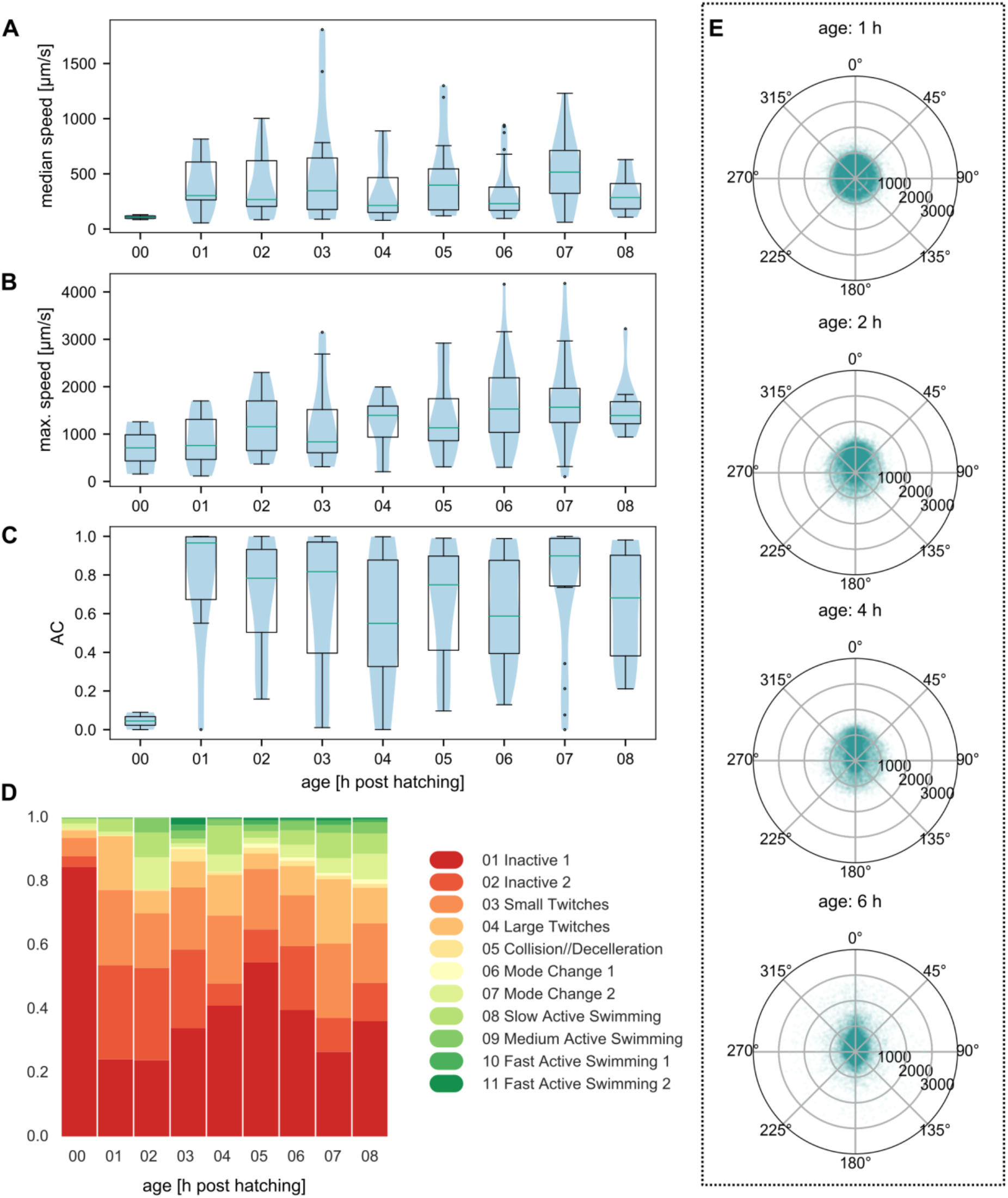
Age related changes in behavioural parameters. (A) Median speeds, (B) maximum speeds and (C) Activity coefficient (AC) for WT animals reared at 14°C at different age post hatching. (D) Distribution of behavioural modes in animals of different age. (E) Speed-turn plots for ages 1, 2, 4 and 6 h post hatching. (N(00)=2, N(01)=8, N(02)=14, N(03)=12, N(04)=15, N(05)=12, N(06)=29,N(07)=18, N(08)=10, number of points per polar plot is 50000)

### Rearing temperature effects on a developmentally regulated behaviour

We (Fig 4) and others have shown that as the *C. intestinalis* larvae go through the post-hatching motile phase of their life cycle they change their behavioural responses at multiple levels. Such behavioural changes are thought to be tightly linked to developmental changes taking place in the larvae. An interesting question that arises is whether this developmental regulation of locomotor and sensory behaviours is robust to different environmental circumstances, possibly through a mechanism of canalization (80-83) or whether it shows plasticity (84-86). Temperature is one key physical parameter that has been shown to affect the speed of most biological processes, acting as a major environmental factor influencing the rate of animal development (87-89) and behaviour(90, 91). The two Ciona species *(Ciona intestinalis* and *Ciona robusta)* (92)occupy a very large part of the world’s coastline from high to low latitudes(93) and they show great adaptability to a range of temperatures. Published studies have used 18°C as rearing and assay temperature for *C. intestinalis* and *Ciona robusta* larvae. However, our local animals belong to the *C. intestinalis* species and develop best at lower rearing temperatures, possibly due to an adaptation to the lower water temperatures in the Norwegian Fjords. We tested whether the lower rearing temperature of 14°C affected the onset of the light-off response that has previously been described by Nakagawa et al.(28). The increase in swimming speed immediately after a light stimulus is considered a hallmark of the older larva that will in its later age seek to settle utilising negative phototaxis. In animals reared at 18°C, the first notable response to a light-off signal was detected at 4 h post fertilisation, coinciding with a reduced average speed of the animals in absence of the stimulus. The sensitivity of this response has been shown to peak in the green part of the spectrum (28), so we tested for light-off response in our animals using a green light (515-530 nm). We calculated the change of speed (i.e. Δ speed (Off), Fig 5B) between the last 10 s during a 1 min light stimulus and the first 2.5 s after the light was turned off (after a 0.5 s latency period). In Fig 5C we present the distribution of the different behavioural modes for the same periods at the end of the light stimulus (L) compared to the onset of the dark (D) for animals at three different post-hatching ages. At 2 and even 4 h post hatching the animals reared at 14°C did not show a significant light-off reaction, with the first notable response observed only at 6 h. The response was notable both as a slight general increase in speed (Fig 5B) and more specifically as higher representation of twitching and medium speed swimming at the expense of inactive modes (Fig 5C). It appears that *C. intestinalis* has the potential to be a great genetically tractable model to answer the question of developmental stability of behaviour.

**Fig 5.**
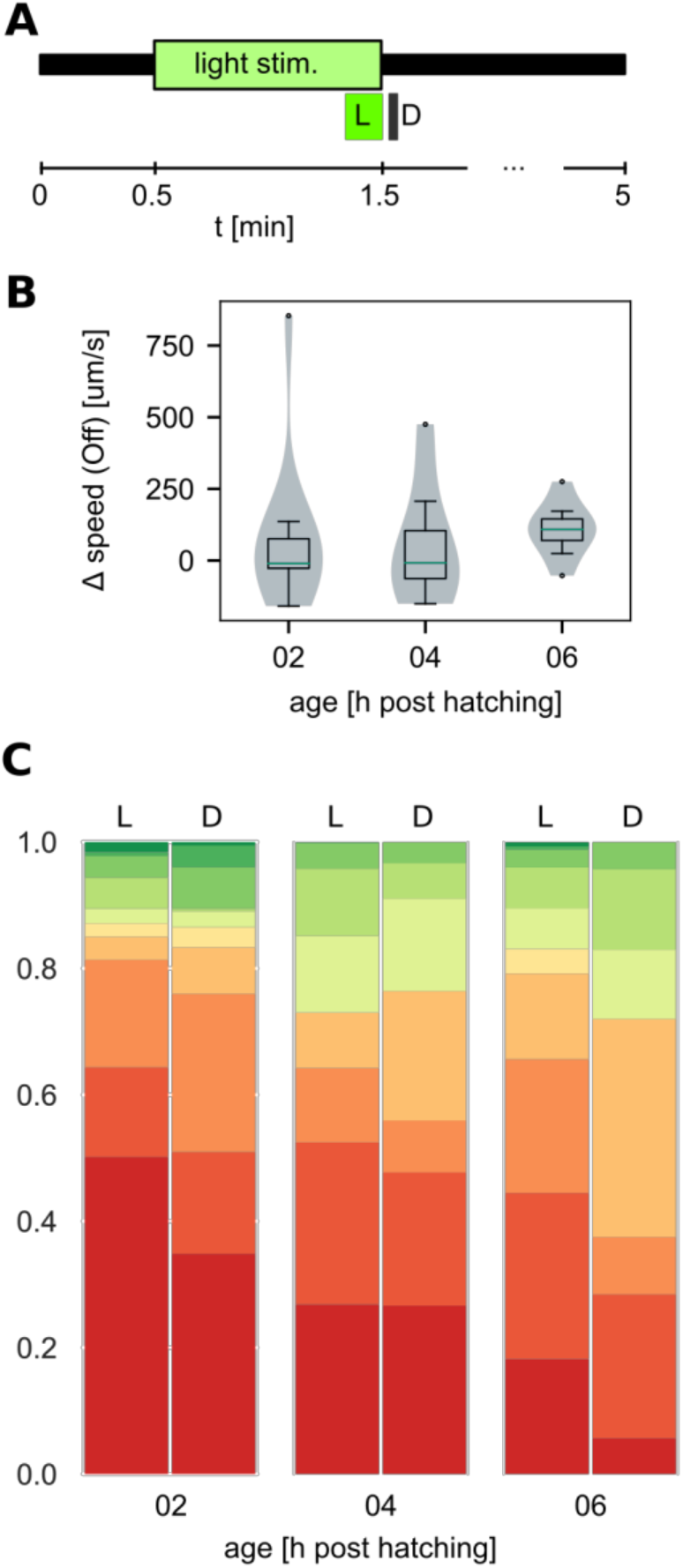
Light-Off response in animals reared and recorded at 14°C. (A) Experimental set-up. Light stimulus lasted 1 minute and the average speed of the animal in the last 10 s of the stimulation (L) was compared to the expected peak of speed upon onset of darkness (D) measured as average speed over 2.5 s starting from 0.5 s after light-off (B) Change in speed between L and D for different aged animals. (C) Distribution of behavioural modes in the last 10 s during a 1 min light stimulus – L and the onset of darkness – D for different aged animals (N2=11, N4=7, Ns=8)

### Rearing temperature effects on behaviour

Animals reared at 18°C had a much narrower time window after hatching in which we could observe active swimming behaviour, with the majority of animals being highly inactive by age 4 h post hatching (data in S2C). We therefore only compared animals of age 0-3 h reared at 18°C to the 14°C reared animals of ages 2-8 h, since we assumed they correspond to the same post hatching development stages. Even when comparing what we assumed to be animals at a similar developmental stage, animals reared and recorded at 18°C still exhibited some differences compared to the ones at 14°C (Fig 6). Their traces were similar in median speed values (Fig 6A) but reach significantly higher maximum speeds (Fig 6B, median at 1864 μm/s for animals reared at 18°C vs. 1440 μm/s for animals reared at 14°C, *p* >0.001). The difference in AC and path complexity is not statistically significant (Fig 6 C, D), but there was a slight but significant effect on thigmotaxis (Fig 6E,

**Fig 6.**
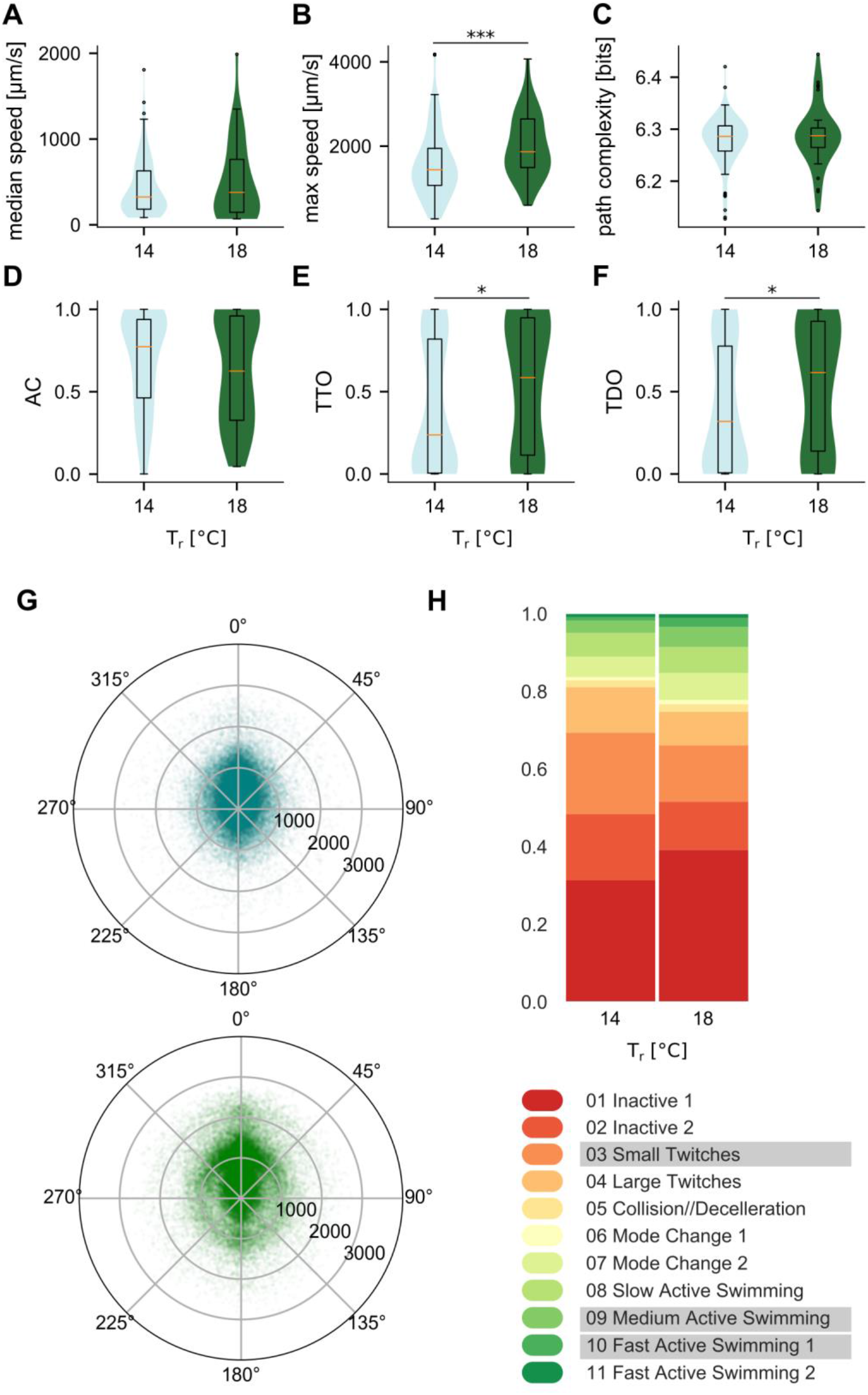
Effect of rearing temperature on behaviour. Rearing temperature effects on (A) median speed, (B) maximum speed, path complexity (C), activity coefficient (D) and thigmotaxis measures (E) TTO and (F) TDO. (G) Polar scatterplots of filtered speed values vs turn values for animals reared and recorded at 14°C (in teal) and at 18°C (in green). (H) Distribution of behavioural modes for the two groups. The modes underlined in the legend are the biggest contributors to the Chi^2^ statistics (data in S1 Table) (N(14°C) = 101, N(18°C) = 36; number of points per polar plot is 100000)

F). The median TTO for animals reared at 18°C was 0.585 vs 0.237 at 14°C (Fig 6E, *p* = 0.035) and the median TDO at 18°C was 0.616 vs 0.319 at 14°C (Fig 6F, *p* = 0.038). At 18°C we also observed a higher representation of medium-high speeds (around 1000-1500 μm/s) in combination with a wider range of turn values, while at lower speeds the variability of turns was smaller (Fig 6G). This was matched with a lower representation of twitching modes and more occurrences of the modes representing swimming at medium speeds (Fig 6H).

### Dechorionation effects

The eggs of *C. intestinalis* are nested in a chorion surrounded by follicle cells (Fig 7A) and the normal development of left-right asymmetry in the embryo has been shown to be disrupted by dechorionation (94). However, transient transgenesis of *C. intestinalis* is predominantly achieved by electroporations, which do require the dechorionation of eggs. We therefore set out to test if dechorionation has specific effects on behaviour. Being aware of any potential effects of dechorionation will be vital for the future interpretation of behavioural phenotypes in transgenic animals. The dechorionated larvae achieved a higher median (Fig 7B, 477 μm/s median vs 324 μm/s in chorionated animals, p=0.0125) and maximum speeds (Fig 7C, 1784 μm/s vs 1440 μm/s for chorionated animals, p=0.0024). The differences in AC and path complexities were not statistically significant in our set (Fig 7 D, E). There was however a slight but significant effect on thigmotaxis (Fig 7 F, G), resulting in a higher median TTO (0.56 vs 0.23 for chorionated animals, p= 0.035) and TDO (0.58 vs 0.32 for chorionated animals, p= 0.033) values for dechorionated animals. The differences in distribution of turns and speeds were less apparent compared to the effect of temperature, but the increased representation of higher swimming speeds in dechorionated animals was evident in the polar scatterplots (Fig 7H) as well as form the distribution of the behavioural modes (Fig 7I).

**Fig 7.**
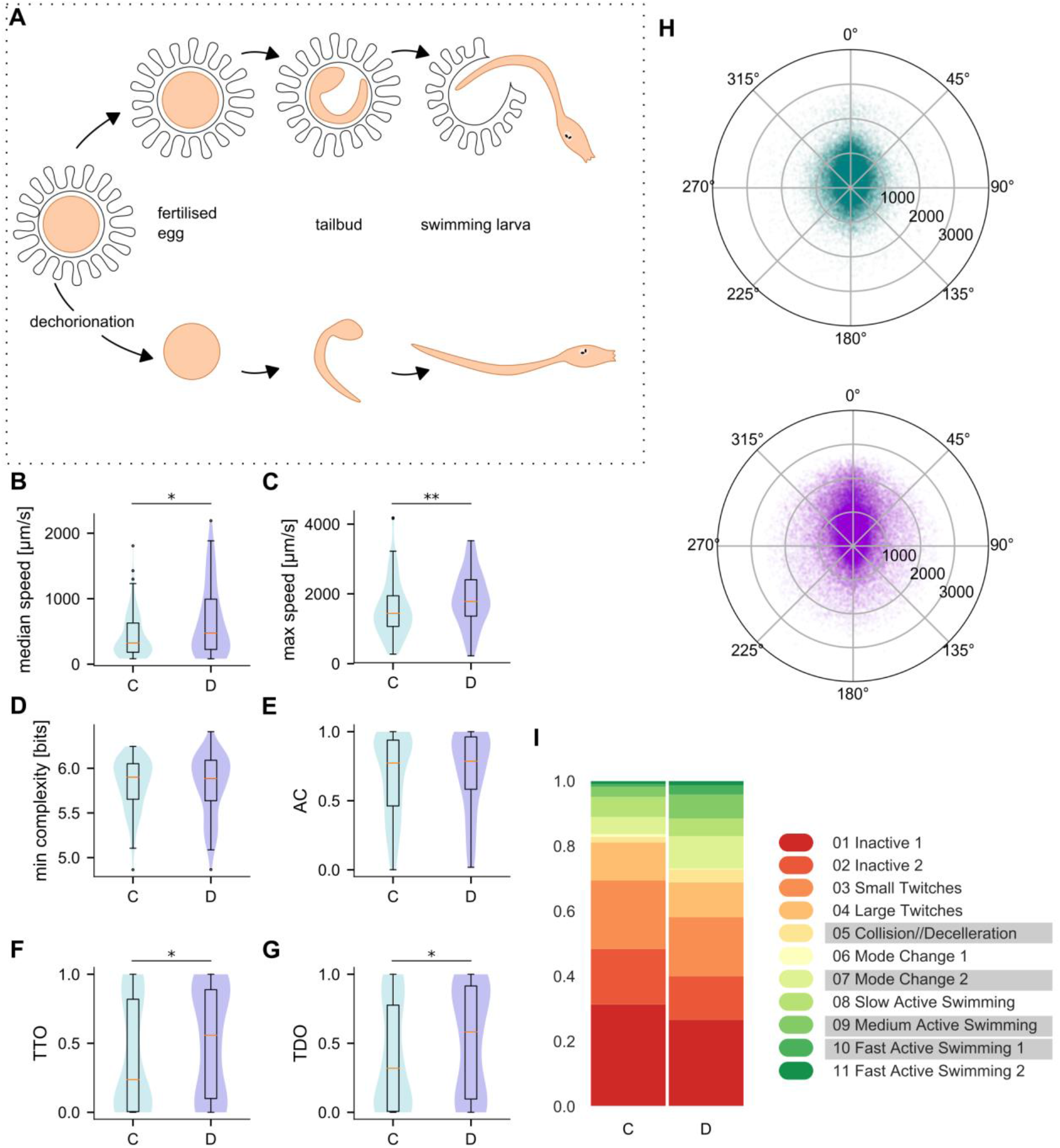
Effect of dechorionation on behaviour. (A) Schematic representation of the *C. intestinalis* embryo development inside the chorion (top) in untreated animals compared to dechorionation and subsequent development of a dechorionated embryo (bottom). We present dechorionation effects on (B) median speed, (C) maximum speed, path complexity (D), activity coefficient (E) and thigmotaxis measures f) TTO and (G) TDO. Chorionated animals are denoted as C, dechorionated as D. (H) Polar scatterplots of filtered speed values vs turn values for dechorionated (purple) animals compared to chorionated animals (teal). (I) Distribution of behavioural modes for the two groups. The modes underlined in the legend are the biggest contributors to the Chi^2^ statistics (data in S1 Table) (N(C) = 101, N(D) = 74; number of points per polar plot is 100000)

### Thigmotaxis and effect of modafinil

Finally, we attempted to influence thigmotaxis in our animals by exposing them to an anxiotropic drug modafinil. In this set of experiments, we compared two groups of animals swimming in 20 mg/l and 2 mg/l solution of modafinil respectively to a control group in DMSO and to the untreated WT set (Fig8, S4). We determined the dosage based on preliminary tests and previously published literature on modafinil effects on larval zebrafish (95). The effect of 20 mg/l modafinil on thigmotaxis was very pronounced and statistically significant (Fig 8E, F). While the median TTO value for the DMSO control was 0,427, it reached 0,669 in the 20 mg/l modafinil group signifying longer periods spent in the outer zone of the arena (p =0.0015). Similarly, the TDO measure shows the modafinil-affected animals travelled much bigger proportions of their total distance in the outer zone of the arena (median TDO at 0.628 in the 20mg/l modafinil group compared to the DMSO control at 0.395, p<0.001). For the 2 mg/l group the distribution for both thigmotactic measures was similar to the WT state and was not statistically different from either of the controls.

The animals affected by modafinil also exhibited an overall more active set of behaviours with much higher representation of the active-swimming modes and less time spent inactively (Fig 8G, H). This resulted in increased median and maximum speeds (Fig 8A, B) and higher AC values (Fig 8D) for animals in 20 mg/l modafinil. Path complexity (Fig 8C) showed fewer differences with the only significant change being the higher minimal complexity values for 20 mg/l group (5.99 compared to 5.85 in DMSO and 5.78 in 2mg/l).

**Fig 8.**
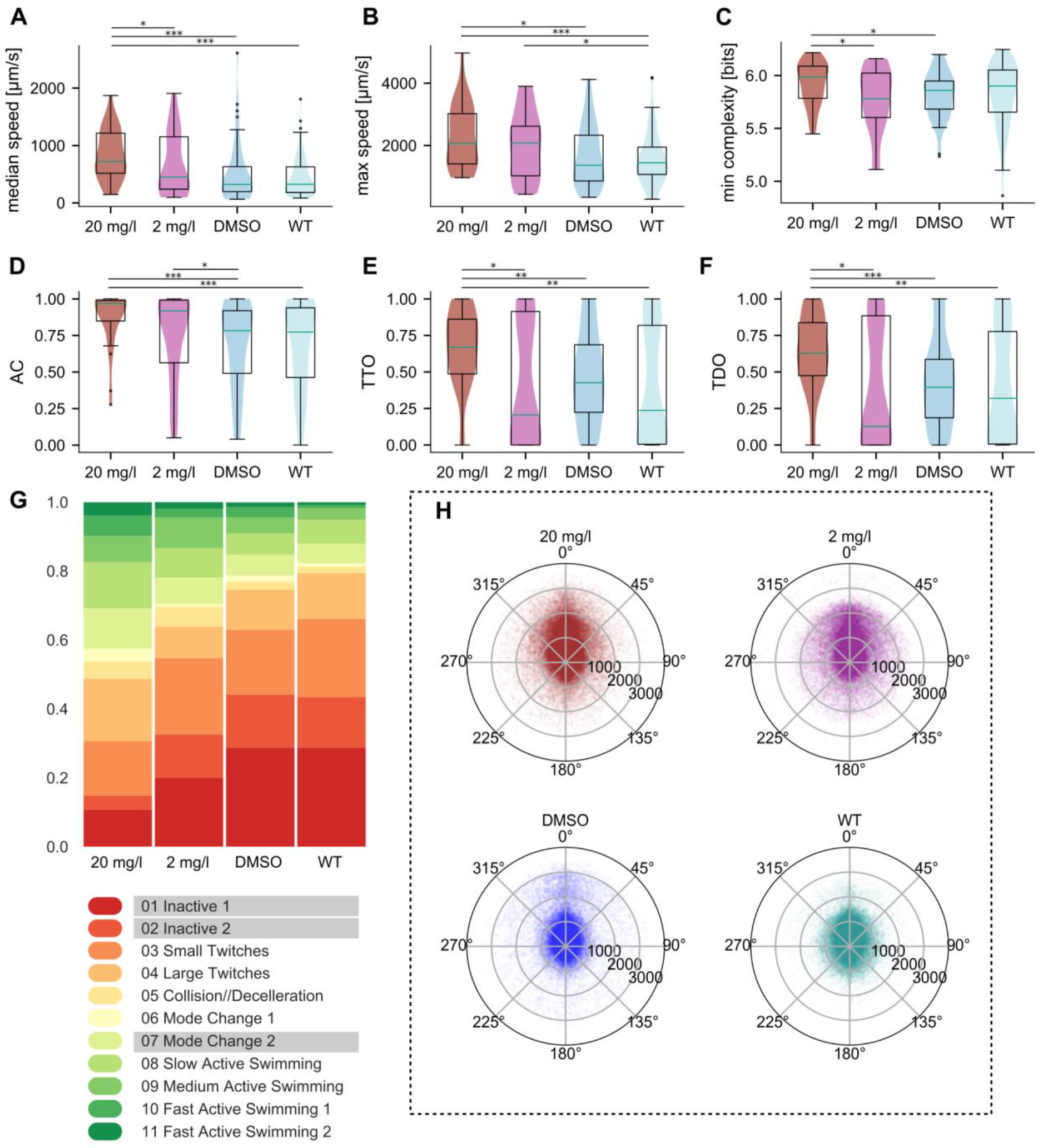
Effect of modafinil on behaviour. Modafinil effects on (A) median speed, (B) maximum speed, (C) minimal path complexity, (D) activity coefficient and thigmotaxis measures (E) TTO and (F) TDO. The two groups affected by 20 mg/l and 2 mg/l modafinil are plotted in dark red and purple respectively, the control group in 0.1% DMSO in blue and wild type animals in light teal. (G) Distribution of behavioural modes for the groups. (H) Polar scatterplots of filtered speed values vs turn values the different groups. (N(20mg/l) = 28, N(20mg/l)= 27, N(DMSO) = 48, N(WT)= 101; number of points per polar plot is 100000)

## Discussion

The chordate *Ciona intestinalis* in its larval form is emerging as a promising organism for neuroethological studies. The present study provides a quantitative description of larval behaviour in different contexts, using biologically relevant features. We performed unsupervised clustering of our data and identify clusters, with which we generate a behavioural ontology. We uncovered some of the behavioural effects of post-embryonic development and dechorionation on larval behaviour and pinpoint the behavioural consequences of crowd size and sensory arousal. Furthermore, we provide evidence that *C. intestinalis* larvae exhibit thigmotactic behaviour that can be modulated by the drug modafinil.

### Quantitative description of *Ciona intestinalis* larval behaviour

The potential of the *C. intestinalis* larva as an organism to perform neuroethological studies has been noticed for several decades. There have been several efforts to perform behavioural studies with increasing sophistication over the years (26-33). However, automated quantitative analysis of *C. intestinalis* behaviour had been hindered by the lack of suitable open-source software with the ability to follow the larvae providing precise positional information over long time series, with few or no identity switches in case multiple animals are tracked simultaneously. In this study, we identified Toxtrac (43) as a suitable open-sourced tracking software, we built customizable hardware and developed an automated behaviour analysispipeline for *C. intestinalis* larvae.

Our analysis suggests that *C. intestinalis* larvae show a surprising amount of complexity in their spontaneous swimming behaviour. Notably, the larvae exhibited a large range of swimming patterns showing significant variation in path complexity (Fig 1F), and individuals used a broad range of behavioural mode sequences (e.g. Fig 3F). The advent of tracking methods has revealed the presence of multiple characteristic scales of organisation in single and multiple animal traces that can be explained only if we consider theoretical frameworks for mobility that extend beyond simple diffusion mechanisms (96, 97). In the future it would be interesting to study the temporal structure of spontaneous swimming in *C. intestinalis* larva in greater detail. Intriguing topics to investigate are for example whether some of the swimming patterns of *C. intestinalis* exhibit a Lévy-like behaviour(49, 98) and whether they may play a role in a dispersal strategy(99, 100). In accordance with our expectations, external sensory cues seemed to influence the swimming strategy of the larvae. Interestingly, path complexity appeared to be modulated in opposite ways by sensory arousal and crowd size (Fig 9).

**Fig 9.**
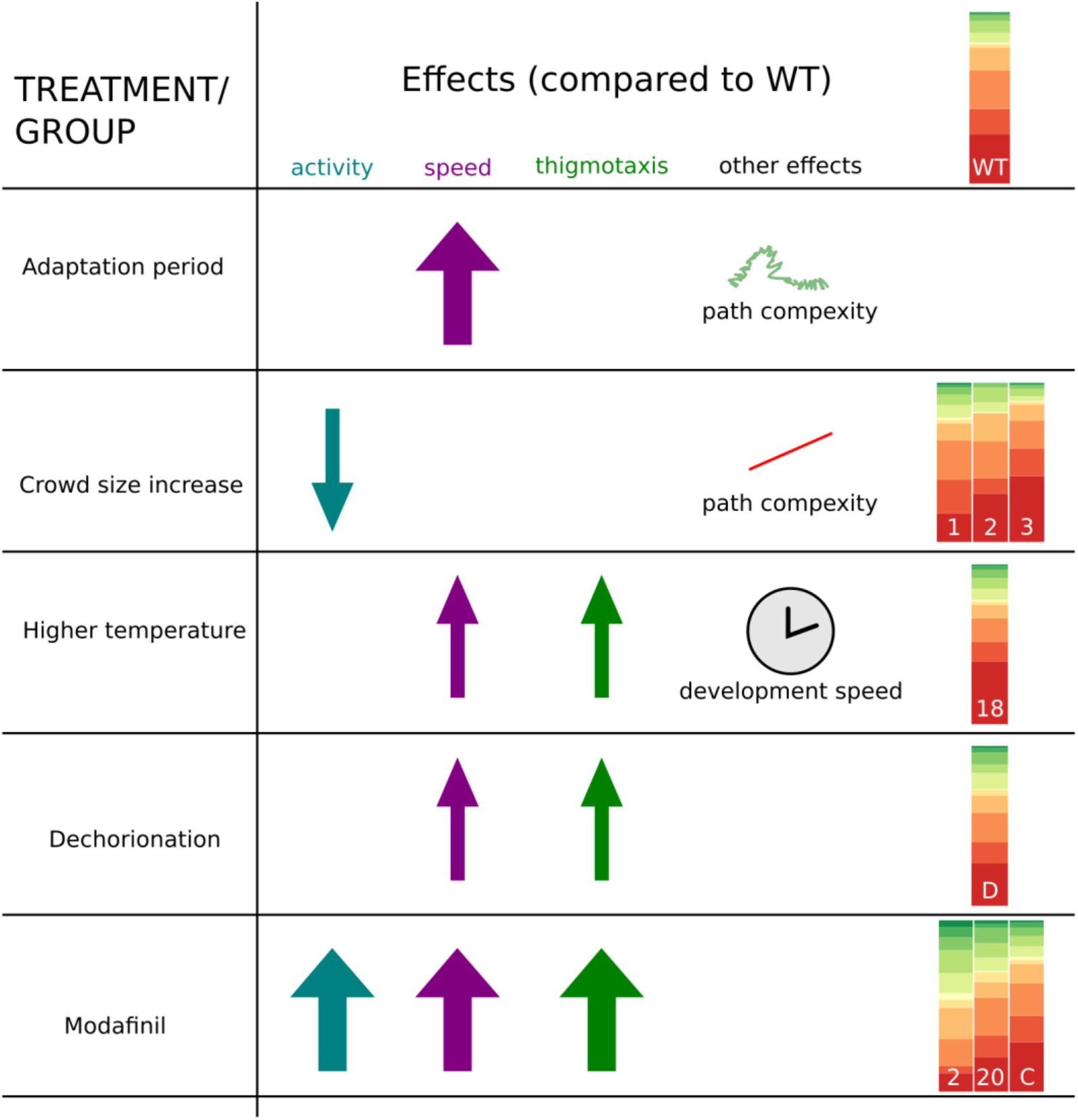
Summary. During the adaptation period (A) *C. intestinalis* larvae exhibited sensory arousal, which translated to higher speeds and increased path complexity. The presence of conspecifics in the arena (B), resulted in reduced locomotor activity, reduced path complexity and a change in the distribution of behavioural modes. We tested the robustness of behaviour in the context of rearing temperature (C) and dechorionation (D) treatments. Finally, the anxiotropic drug Modafinil (E) was able to modulate thigmotaxis, arousal and the overall state of animal activity, by changing the distribution of the behavioural modes.

The sensitivity of our measurements allowed us to quantify the activity levels of the animals in different experimental contexts. We found that larvae showed both bursts of activity and bouts in irregular intervals. Intermittent locomotion(101) is one of the various morphological (102)and behavioural strategies(103, 104), often under evolutionary pressure(105), that have been implemented by moving organisms, including marine larvae in order to conserve energy(106). An alternative hypothesis is that fluctuations in activity levels may reflect an adjustment of motility to small changes in flow taking place in the arena. Similar bouts and bursts of activity characterise zooplankton motility in turbulent flow(107).

Quantitative behavioural analysis has been moving from subjective observation, and imprecise annotation of behavioural data, towards the automated recognition and classification of behaviours amongst very large data sets (2, 4, 7, 108). Some of the breakthroughs that have permitted this progress have focused on generating low-level representations of behaviour for automated analysis and automated classification algorithms of behaviours. The ultimate goal is to break down complex behaviours into their constituent building blocks. In this study, we employed a clustering methodology, using first unsupervised clustering of a minimal feature-set to identify behavioural modes, followed by training a K-Nearest-Neighbours classifier in order to classify all of our data corresponding to approximately 1.8 million observations from 850 data-frames. We found 11 distinct clusters that were classified in an equal number of basic behavioural modes. These 11 behavioural modes provide an unbiased way to dissect the structure of behaviour and will allow us to systematically classify complex behavioural phenotypes that result from pharmacological, genetic or optogenetic perturbations (109-112). However, our current automated image based tracking approach is relying on marking each animal with a centroid rather than segmenting out the entire shape of the animal in order to generate an outline or a skeleton. We therefore are lacking postural information that would enrich our dataset significantly. This presents an important next step towards obtaining a complete ethogram of *C. intestinalis* larval behaviour. We note however, that clustering data points from centroid analysis already allowed us to describe a number of behavioural modes accurately. Furthermore, we are testing our animals in an open field arena that is suitable for recording a relatively small number of animals, possibly in a setting that is relatively distant to the natural ecological niche of the larvae. This problem is faced by numerous experimenters who are trying to obtain high quality tracking data in a controlled environment(5). We envision that in the future the use of larger arenas and the ability to deliver multiple sensory stimuli reaping the benefits of the open architecture of the behavioural setup, will allow us to study other ecologically relevant behaviours such as settlement behaviour and metamorphosis more closely.

### Arousal from transfer to new environment

Animals have the ability to modulate their readiness to react to sensory cues, in a phenomenon known as arousal. This modulation is very obvious when comparing the states of sleep and wakefulness. In addition, during the awake period, animals are able to enter short-term behavioural states, during which they exhibit heightened activity and general or specific sensory stimulus responsiveness and thus are able to anticipate and address sudden challenges (113). Here we report that *C. intestinalis* larvae are in a state of arousal during the first minutes of being placed in the arena (Fig 2, 9). Generalised arousal is thought to be widespread across vertebrates (59, 60) but the detailed neuronal and molecular mechanisms are still poorly understood. In fact, amongst invertebrates, there is evidence that sensory arousal is present in Aplysia(114) and in ecdysozoans, like the nematode worm *C. elegans* (62, 115) and *Drosophila melanogaster(116,* 117). In the case of *C. elegans,* sensory circuits involved in sensing high threshold mechanical and noxious stimuli are implicated in the heightened state of arousal. Given that the arousal exhibited by the *C. intestinalis* larvae is likely due to mechanical stimulation from transfer to the arena it would be interesting to study the contribution of mechanosensory circuits to this behavioural state. In the case of Drosophila, acute sensory arousal is more apparent when comparing between states of wakefulness and sleep(118). A first clue as to which circuits might be involved comes from the observation that arousal in *C. intestinalis* is modulated by the drug modafinil (Fig.9). Modafinil has been classified as a psychostimulant and has been extensively used in narcoleptic patients in order to address sleep related disorders. It is thought to act as a selective dopamine (119) and norepinephrine transporter inhibitor (120), thus raising the possibility that monoamine signalling plays an important role in modulate the arousal state in *C. intestinalis* larvae. In this study we show that beyond modulating the arousal state of the animals, modafinil appears to alter the activity coefficient. The larvae also showed less quiescent periods with a higher activity coefficient. This has also been observed in mice and zebrafish. In mice, modafinil results in wake-promoting action, possibly via dopamine transporters(121), while in zebrafish modafinil shortens the periods of sleep(122).

### Age related changes

A large number of animals exhibit behavioural changes linked to post-embryonic development of their nervous system. For example, in *Xenopus laevis,* locomotor activity patterns are modified as the animals transition from sessile hatchlings to free-swimming larvae(76), through changes in the cell properties of neurons(123) and a nitrogen oxide signalling mechanism (75). Other examples of post-embryonic changes in behaviour include the photoresponses of the stick insect *Carausius morosus* (77) and the chemotactic responses in *C. elegans(124). C. intestinalis* larvae are also subject to post-embryonic developmental changes and associated behavioural modifications. It has previously been reported that light and gravity dependent behaviours change during the larval life (26, 31). Here we quantified how progression through larval life changes behaviour of the animals, and found age dependent differences in the distribution of behavioural modes (Fig 4). This puts a quantitative angle to previously reported findings in the literature(30). What may be the mechanisms that bring about these changes in behavioural modes used in larvae of different ages? One possibility is that some neurons of the larval system fully differentiate and connect to the nervous system only after hatching. Indeed, there has been evidence for post embryonic terminal differentiation of dopaminergic cells in *C. intestinalis* larvae (125). The authors of the study postulated that dopamine might modulate the neural circuits involved in the age-dependent changes in swimming behaviour of the larva. The genetically tractable *C. intestinalis* is proving to be a powerful model for providing in-depth insight into developmental processes in the post-embryonic nervous system and the functional basis of locomotor dynamics changes throughout larval life. Notably, the use of a fast-growing larval animal with associated changes in body proportions and shape, provides us with the opportunity to understand how alterations in biometrics and sensory capacity may relate to simultaneous changes in locomotory behaviour.

### Crowd size effects on larval behaviour

Many behaviours, such as mating, shoaling, schooling, aggressive encounters rely on the interaction of two or more individuals. These interactions largely depend on achieving a coordinated movement between individuals and the entire group(126).

Previous work in *C. intestinalis* has shown that larvae can aggregate into a column when placed in a three dimensional chamber and that they can form swarms, especially upon agitation of the water(29). Also it has been shown that larval distribution can change in the presence or absence of light(30). Notably, it has been reported that ascidian behaviours prior to settlement are largely influenced by conspecifics, while the larvae exhibit a form gregariousness(35, 127). However, these are largely qualitative observations that were made in the course of experiments that were not designed to specifically address the interactions between conspecifics. Further difficulties for providing a quantitative description of larval interactions stem from the lack (until recently) of automated tracking software that faithfully maintained the identity of each tracked animal(5). Taking advantage of the ability of ToxTrac to maintain the identity of multiple animals in the same arena we attempted to determine whether there are differences when single vs multiple individuals are placed in the arena. We find that the presence of two or three animals in the arena can already result in a few changes in the measured behavioural parameters (Fig 9). Notably, a significant change in one of the thigmotactic indices is also observed (Fig 3D). Interestingly, an enhancement of thigmotaxis in individual versus group context has been observed in the case of ants(128). Given the past literature it will be interesting to determine if and how *C. intestinalis* larvae achieve coordinated movement. It is believed that the type of distributed sensing required to generate robust collective behaviour is rather simple, requires rudimentary circuits and thus it may be widespread across different animal taxa(129). *C. intestinalis* has a small nervous system and thus is ideal to study the neural circuits controlling pairwise and group-level behaviours. We note that a limitation of our method is that our crowd size experiments were conducted in an extremely small volume of sea water compared to the ethologically relevant volumes that these animals would encounter in the sea. Future experiments should be conducted in larger arenas that may be ethologically more relevant for crowd size experiments.

### Behavioural robustness to altered rearing conditions

Temperature is a known modulator of key physiological processes and behaviours in numerous animals(130). For example, thermal rearing conditions can affect the dispersal of adult spiders(131), the host-seeking behaviour of parasitic nematodes(132), mating behaviour in Drosophila(133), and the feeding behaviour in the mud snail *Heleobla australis(134).* To contextualise our data we compared our main wild type group reared and recorded at 14°C to animals at 18°C, since behaviour of *C. intestinalis* larvae has previously often been studied at 18°C e.g. (27, 31, 32) or even at room temperature (29). We describe the distinct difference in the speed of post hatching development at different rearing temperatures (Fig 4, S2) At 18°C the period of higher spontaneous locomotor activity coinciding with the lack of significant responses to light-off stimuli lasts for ca. 4 h post hatching (28). In animals at 14°C we can detect the first significant, yet still weak, responses to the light-off signal at 6 h post hatching (Fig 5), coinciding approximately with the period of higher locomotor activity in animals up to 8 h post hatching. Apart from this clear influence in the rate of post hatching development, the higher temperature seemed to have little effect on the animals’ behaviour, but we do note the higher representation of medium-high speeds in combination with a wider range of turn values (Fig 6). It may be the case that rearing temperature has no strong behavioural defects in *Ciona intestinalis* larvae, possibly through a buffering mechanism. Alternatively, we may have not identified the behaviours and sensory modalities affected by rearing temperature. This is a plausible explanation in light of significant evidence suggesting that not all sensory modalities are affected equally by the rearing temperature or from deviations from that, since olfaction appears to be particularly strongly affected (135-139) compared to other sensory modalities in other organisms.

Yet another treatment that can challenge the behavioural robustness of *Ciona intestinalis* larvae is the enzymatic removal of the chorion that envelopes the eggs, a process termed dechorionation.

This enzymatic treatment is an essential step in the generation of transgenics via electroporation. However, it can potentially interfere with the establishment of brain asymmetry in the ascidian brain, which is dependent on an intact chorion(140). Given that left-right asymmetries in behaviour and in nervous system structure are abundant phenomena across different animal taxa(141), it was of paramount importance to understand, in the first instance, the effects of dechorionation to spontaneous larval swimming. The behavioural comparison of chorionated versus dechorionated animals revealed differences in speed and thigmotaxis. Unexpectedly, dechorionated animals swam faster and showed higher thigmotaxis levels (Fig 10D). These observations suggest that future quantitative behavioural studies making use of electroporated transgenic animals need to use dechorionated animals for ‘wild-type’ controls rather than larvae hatched from chorionated eggs. Nonetheless, it is encouraging to see that dechorionated larvae are in many behavioural parameters indistinguishable from chorionated egg derived larvae.

### Thigmotaxis and modafinil effects

Using an open-arena to monitor our animals, we noticed that a large fraction of larvae exhibited strong thigmotactic behaviour. This appears to be an adaptive behaviour that has been observed in numerous organisms, where the circular wall of the arena allows the animals to exhibit a defensive response (i.e. to hide from potential predators) and facilitates their orientation in space(142). Therefore, it is not unlikely that thigmotaxis presents an evolutionarily conserved behaviour. One may wonder what role thigmotaxis plays in the larvae of *C. intestinalis.* In fact, almost thirty years ago it was hypothesised that thigmotaxis, amongst other behaviours, may be involved in the selection of habitats for larval settlement(35). Interestingly, we have been able to demonstrate that physiological (rearing temperature) and morphological changes (dechorionation) can affect the animal’s ability to perform thigmotaxis. Moreover, we found that modafinil increased thigmotaxis levels in *C. intestinalis* larvae. This is interesting in light of the fact that the effects of modafinil treatment in both humans and other animals has shown variable effects. In some cases, it acts as an anxiogenic drug like in one study in humans (143)and in others as an anxiolytic drug such as in marmosets(144). Notably, one study showed that modafinil increased the exploratory behaviour of mice in a dose dependent manner (145). It has also been shown that Modafinil can reduce thigmotaxis levels in zebrafish, (95). The strong effect that the anxiotropic drug modafinil has on *C. intestinalis* larval thigmotaxis is evidence that a common mechanism might mediate thigmotaxis across taxa(63, 95, 146). Future work should explore the molecular and cellular underpinnings of thigmotactic behaviour in *C. intestinalis* larvae and aim to understand the ecological context in which it may be used.

### *Ciona intestinalis* as a neuroethological model

For years, cutting edge functional imaging and automated behavioural analysis was the privilege of a few model organisms. However, there has been a turn of tide in recent years. For one, the understanding that comparative studies are very important in order to draw conclusions on the generality of biological phenomena and mechanisms, drew the attention to non-model organisms. At the same time, functional imaging techniques and molecular toolkits started to became highly adaptable, in order to meet the requirements of previously intractable nervous systems(147). The same trend was observed with open-source machine vision and machine learning tools that could be used to automatically track a large variety of organisms with different shapes and behavioural repertoire (16, 18, 148). The larval form of *C. intestinalis* fulfils several criteria that would allow it to ride this wave of change in neuroethology.

Phylogenetically, *Ciona intestinalis* is positioned at a key evolutionary node, as a member of the sister group of vertebrates(149). Its compact nervous system showcases numerous developmental mechanisms and gene regulatory networks common to all chordates, including vertebrates(150). The complete mapping of the larval wiring diagram using Electron Microscopy (22-24) allows for structural framework upon which to generate testable hypotheses. Most importantly the genetic and genomic toolkit (41, 151) together with the ease of transgenesis and the transparency of the larva make this organism particularly amenable to optogenetics and functional imaging. Notably, the Genetically Encoded Calcium Indicator (GECI) GCaMP6s, has already been used in *C. intestinalis* to study developmental calcium dynamics (152-154). The adoption of GECIs in *C. intestinalis* combined with our approach of automated behaviour recognition and analysis have allowed us to systematically dissect the larval behavioural repertoire and image the entire nervous system in vivo, with the ultimate aim to elucidate the neural networks underlying behaviour. With our approach, we were able to show that we can quantify larval behaviours automatically and identify novel behaviours (thigmotaxis) and behavioural states (arousal). This approach also allowed us to investigate the robustness of the behavioural repertoire under diverse environmental, developmental and pharmacological conditions. Future work, includes obtaining a more detailed mechanistic understanding of the stimulus driven behaviours, social interactions and learning paradigms.

## Methods

### Animals

Adult Ciona intestinalis were collected locally from the Bergen area and Sotra Island, Norway. We incubated them in filtered seawater at 10°C under constant illumination to stimulate egg production. Eggs and sperm were obtained from individual animals to perform in vitro fertilisation. Part of the eggs were dechorionated using Na-Thioglycolate and mechanical dechorionation(155). Both eggs with and without chorion were fertilized at the same time and incubated in artificial sea water (ASW, Red Sea Salt) at either 14 or 18 °C. The post hatching age of animals is referred to relative to the onset of hatching of larvae from the chorion.

### Set-up for behavioural experiments

Animal behaviour was recorded in a custom-made setup developed in our lab (Fig 1A). Using a 3D-printed PLA mould, we made single-use agarose arenas (0.8% in ASW, by Invitrogen, USA,). The arena was nested inside a PLA ring with infrared (IR, peak emission 850 nm) LEDs, which provided dark-field illumination of the animals without stimulating their photoreceptors. The ring also held a small thermometer (DS18B20, Maxim Integrated) positioned close to the arena and was placed on top a Peltier element with a thin layer of ASW underneath the agarose to improve heat conduction and image quality. Light stimulation was performed using LED illumination (green LED in NeoPixel LED array; emission 515–530nm) and an IR filter (cut-of at 780 nm) positioned in front of the camera. Videos were recorded using an IR sensitive monochrome camera (DMK 33UP1300, The Imaging Source, Germany) and IC Capture software. An Arduino based circuit, interfacing with a GUI written in Python, provided stimuli and PID-controlled temperature control.

### Recordings

1–3 animals were placed in an agarose arena (10mm in diameter and 3mm high, approximate volume 236 mm^3^). Each animal in behavioural experiments was first filmed for a period of 15 minutes during acclimatization to the arena (at 10 frames/s). Subsequently 1 to 3 videos of 5 min duration were filmed at 30 frames/s to analyse either base line behaviour or effect of stimulation, rearing temperature or added drugs on behaviour. For modafinil experiments the animals were first transferred to a dish containing DMSO or modafinil and then immediately transferred to the arena which also contained DMSO or modafinil (2 or 20 mg/l) as schematically presented in Fig 8.

### Video conversion and analysis

Videos were analysed using ToxTrac software and custom-made software using OpenCV and python environment (Fig 1C). For each video all frames were enhanced with Contrast Limited Adaptive Histogram Equalization (CLAHE) with a clip limit of 1 and a tile grid size of 50×50 pixels. After histogram equalization noise was reduced with a median blur with a tile grid size of 5×5 pixels. To input bright-background videos into the ToxTrac software, all frames were inverted by subtracting from a true white frame of equal size. Within the ToxTrac software, the ID algorithm used in our study was 2TCM sel. by Hist (MEE).

### Data analysis and statistics

All data analysis was performed with python using the numpy, pandas, scipy, scikit-learn and matplotlib libraries. For every analysed video the position of the centre of the arena is determined with a Hough Circle Transform algorithm in OpenCV in python. For every trace the [x,y]-positions are corrected so that [0,0] was at the centre of the arena. All positions were then multiplied by the factor of 11.56μm/pixel for the setup the recording originates from. From these positions distances, speeds and subsequently all other parameters are derived. We excluded animals that were completely immobile and hence indistinguishable from dead form further analysis by excluding all traces where the maximal displacement from the starting position was less than one body-length (comprising approximately 10% of all examined traces). Similarly, traces where the animal was tracked for less than 2000 frames were considered unrepresentative and excluded from further analysis.

The tracked centre-point corresponded predominantly to the animal’s head. To filter out noise caused by the undulatory movement of the head during swimming the speed sequence values were filtered with a 1 Hz low-pass filter. All speed values presented are therefore filtered speeds and turn values refer to values calculated between coordinates 5 time-points apart for the same reason. Whenever statistical significance was tested we used nonparametric test for all non-normally distributed data, namely Levene’s test for equality of variances, Kruskal-Wallis analysis of variance and Mann–Whitney U test. To compare the similarity of distributions in the scatterplots of speed vs turn values presented in this paper we compare the sorted Mahalanobis distances of these distributions (data in S3). Where relevant, *X* statistic was calculated for behavioural modes distributions and the greatest contributor underlined in the figure legend (see Supplemental material for values).

### Clustering of behavioural modes

To identify objective behavioural modes we attempted unsupervised clustering of a minimal feature-set that describes the behaviour of the larvae. The featureset was created as follows: For each good recording the velocity vectors (ρ, ϕ) were calculated from coordinates 5 frames apart. Values p, Ap and Aϕ were used as measures for speed, acceleration and turns respectively. For each point the mean of a sliding window of [-25:+25] frames was used to include information of past and future movement. This results in a dataset of three features and ca. 1.8 million observations. Clusters in the dataset were identified using an agglomerative clustering algorithm with ward-linkage. To determine the optimal number of clusters we identified the point where adding more clusters would not reduce the total distance of all points to their respective cluster centre drastically.

### Classifying

Using the clustered dataset, we trained a K-Nearest-Neighbours classifier that takes a recording expressed in the features mentioned above, and assigns each point in this recording to a cluster. We classified all collected traces with this classifier, and inspected the original videos with the assigned clusters superimposed in order to assess the biological relevance of each cluster. We found that apart from cluster 0, which turned out to be the result of an artefact from data were there was insufficient datapoints in the window for averaging, we could identify distinct behaviours for the remaining eleven clusters. Several of these clusters described the same biological behaviour but resulted in different clusters as a result of the tracking marker being placed in either the head or the neck of the animal by the tracking software (For an illustration of the speed and turn values present in the different clusters see data in S1).

### Data availability

The designs for the behavioural setup components that were 3D printed can be found here: https://github.com/ChatzigeorgiouGroup/Rudolf-Dondorp-2018/tree/master/3d%20Cad%20files The dataset used for this study can be found here: http://doi.org/10.5281/zenodo.1298978 The analysis code is located in our GitHub repository: https://github.com/ChatzigeorgiouGroup/Rudolf-Dondorp-2018/tree/master/Code

## Acknowledgments

We would like to thank Felicia Keulder-Stenevik for animal collection, Mie Wong and Kushal Kolar for comments on the manuscript. M.C. would like to thank Lionel Christiaen, Alberto Stolfi and Florian Razy-Krajka for introducing him to *Ciona intestinalis* experimental methods.

